# Temporal Dynamics of Flexible Cognitive Control

**DOI:** 10.1101/2025.10.03.680146

**Authors:** Chengyuan Wu, Carol A. Seger, Canhuang Luo, Ying Zhou, Jianfeng Jiang, Qi Chen

**Affiliations:** School of Psychology, Shenzhen University, Shenzhen, China; Department of Psychology, Colorado State University, Fort Collins, United States; Department of Psychological and Brain Sciences, University of Iowa, Iowa City, United States

**Author notes:** Address Correspondence to Qi Chen, Address: *School of Psychology, Shenzhen University, No. 3688, Nanhai Avenue, 518060 Shenzhen, China*.

**Keywords:** cognitive control flexibility, dynamic environment, hierarchical Bayesian model, reactive control, proactive control

## Abstract

In dynamic environments, flexible cognitive control adaptively adjusts processing through proactive mechanisms deployed in advance and reactive mechanisms engaged upon conflict. Previous studies have primarily focused on identifying neural networks supporting specific control components, while less is known about how multiple components interact over time to support adaptive control. To characterize these temporal dynamics, we combined EEG recordings with a face-word Stroop paradigm under changing conflict environment. A hierarchical Bayesian model was used to estimate trial-wise learning rate, predicted conflict level, and prediction error, providing computational indices of cognitive control flexibility. Neural correlation analysis revealed that these variables correlated with Theta, Alpha, and Beta oscillations in distinct brain regions. Connectivity analysis among these regions indicated enhanced cross-frequency directional interactions triggered by stimuli. Furthermore, connections reflecting updates to predicted conflict level prior to stimulus onset indexed individual strength in proactive control, while connections reflecting learning rate updates after stimulus onset indexed reactive control. These findings highlight how oscillatory dynamics coordinate multiple control components and provide new insight into how proactive and reactive control emerge as distinct modes within this interconnected neural architecture of flexible cognitive control.

## Introduction

Cognitive control enables individuals to effectively extract task-relevant information and suppress dominant responses during goal-directed behavior [1]. A key feature of cognitive control is its flexibility. The control processes must be capable of dynamically adapting to task demands in a constantly changing environment. Adaptive adjustments in control processes can be implemented through two complementary modes: proactive control, which involves the maintenance of task representations and the anticipatory deployment of control to deal with upcoming cognitive conflicts, and reactive control, which is transiently triggered by stimuli to resolve unexpected cognitive conflicts [2,3].

The flexibility of cognitive control allows the brain to modulate the relative engagement of proactive and reactive control in response to changes in the environment. Factors such as proportion congruency, volatility, the stage of cue presentation, intertrial interval, and working memory load can all influence the trade-off between proactive and reactive control [4–6]. Previous studies have attributed the mechanisms underlying a range of conflict adaptation effects [7–10] to proactive and reactive control, and have attempted to explain these processes using computational models. Associative learning models, typically developed as extensions of conflict monitoring theory [11,12], simulate proactive and reactive control by biasing connection weights or strengthening task representations [13–16], whereas reinforcement learning models implement these control modes by continuously updating the expected value function through temporal-difference learning to determine whether proactive or reactive control should be engaged [17–19].

However, these previous studies tended to deliberately divide proactive and reactive control and assign separate parameters based on their presumed nature, which introduced certain limitations. While such approaches captured some features of cognitive flexibility, they also constrained the efficiency of adaptive adjustments in dynamic environments. To address this limitation, Jiang and colleagues [20,21] proposed a more compatible framework based on hierarchical Bayesian perspective. In this framework, the brain adaptively updates trial-by-trial posterior distributions of latent variables that quantify the conflict context according to previously experienced conflicts. Control is then deployed according to the expected values of these distributions to optimize performance on the upcoming trial. Although the model does not explicitly separate proactive and reactive control modules, both modes emerge from the inference process. This framework can reproduce adaptive effects typically attributed to proactive and reactive control without requiring their pre-defined segregation.

Prior research suggests that the adjustment of control processes is jointly implemented by the dorsal anterior cingulate cortex (dACC) and a network of tightly interconnected and often coactivated cortical and subcortical regions [22–24]. The dACC monitors and evaluates input signals, which are preliminary representations of cognitively meaningful information generated by cortical and subcortical areas, such as value, feedback, emotion, and internal states. Of these regions, the anterior insula (AI) is one of the most representative. The AI shows phasic responses to a range of salient events that may signal the need to adjust control [23,25]. After the dACC integrates this information and specifies control signals, the dorsolateral prefrontal cortex (dlPFC) modulates specific task-relevant processing pathways. In addition, subcortical regions regulate global parameters of information processing, collectively facilitating the execution of control signals[18,24,26].

In dynamic environments, the brain must continuously update its anticipation in response to contextual changes in conflict and execute control processes in both proactive and reactive modes [26]. Studies have shown that these control processes are instantiated through a network comprising the frontoparietal cortex and its connected subcortical regions, particularly the insula and striatum [21,27,28]. Within this network, volatility of control demands is represented by the insula, expected and unexpected conflicts are tracked by the caudate nucleus, and the implementation of proactive and reactive control is reflected in broadly distributed components of the prefrontal network including the anterior cingulate cortex (ACC) and adjoining regions.

However, although it is increasingly recognized that different phases of a task place distinct demands on flexible cognitive control, the temporal dynamics of the neural mechanisms underlying such flexibility remain unclear. Previous research on neural oscillations has provided initial evidence that sustained adjustments of cognitive control are implemented through multiple oscillatory components [29]. These components include frontomedial Theta oscillations which support ongoing monitoring and evaluation of conflict [30–32], Alpha oscillations which contribute to the inhibition of task-irrelevant process [33–35], and Gamma oscillations which facilitate the excitation of task-relevant process [36,37]. However, these studies have often been limited to a single task phase or a single frequency-band component, leaving an incomplete understanding of how different frequency bands associated with distinct cognitive components coordinate across task phases with varying demands. Moreover, if the temporal course of flexible cognitive control is indeed implemented through interactions among neural oscillations at different frequencies, an important question arises: to what extent can proactive and reactive control be reflected as dissociable signatures within this neural process? While behavioral and neuroimaging evidence already supports such a temporal dissociation [38,39], it remains unclear whether oscillatory dynamics provide corresponding neural signatures—a gap the present study seeks to address.

The present study employed a Stroop paradigm under dynamic conflict contexts (Figure 1A and 1B), combined with high-density EEG, to investigate how neural oscillations contribute to flexible cognitive control. We first used hierarchical Bayesian modeling [20,21] to estimate trial-wise prediction error, learning rate, and predicted conflict. Linear mixed-effect models were then applied to test how oscillatory power in distinct frequency bands correlated with each of these variables at specific time points, and parallel source reconstruction analyses were conducted to identify the cortical regions where these neural correlations were centered. After establishing these neural correlations, we used Granger causality modeling to examine how frequency-band components associated with the model variables interacted over time to support control processes. To further determine whether proactive and reactive control could be identified as dissociable signatures emerging from these oscillatory interactions, we introduced the trial-wise correlation between prediction error and response time as an index of individual differences in control mode [21]. Within this framework, we hypothesized that connectivity patterns reflecting the updating of predicted conflict would capture proactive control, as this adjustment process supports anticipation before the onset of conflict. In contrast, connectivity patterns reflecting the updating of the learning rate were expected to capture reactive control, as this process reflects adjustments to environmental volatility after the onset of conflict [2,26]. These targeted hypotheses allowed us to assess whether the temporal dissociation between proactive and reactive control, long posited in the Dual Mechanisms of Control framework [2,4,14], can be observed as emergent signatures in oscillatory dynamics. Together, these analyses clarified the neural architecture that supports adaptive behavior in dynamic environments.

**Figure 1.**
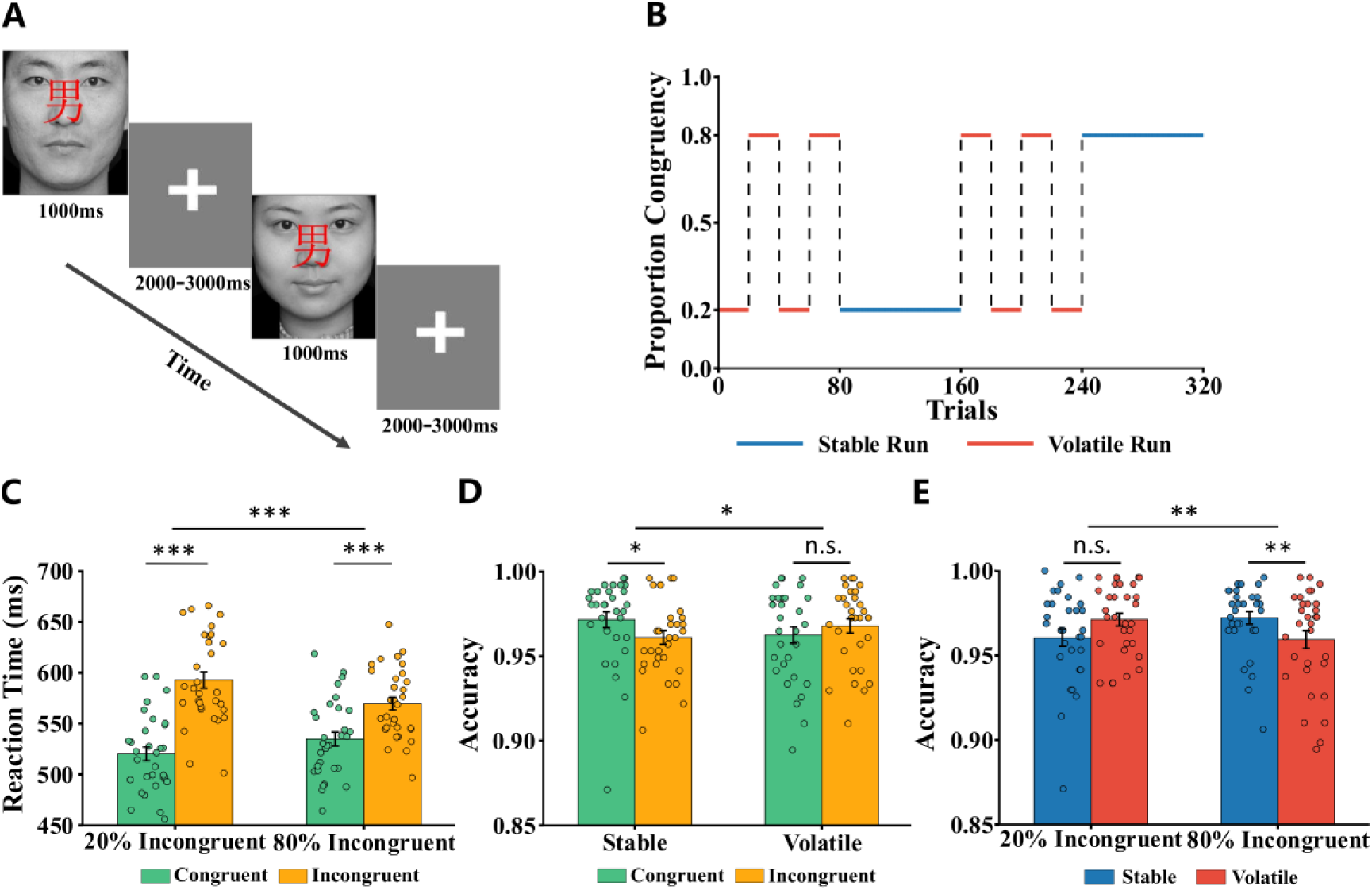
Experimental design and behavioral results. (A) Two sample trials showing the compound stimuli (face with overlaid gender character) and timing of presentation. The first of the two trials illustrates a congruent stimulus (male face paired with the character for “male”), and the second of the two trials illustrates an incongruent stimulus (female face paired with the character for “male”). (B) Proportion of incongruent trials under volatile (red) and stable (blue) runs. (C) Mean RT plotted as a function of current trial congruency and proportion congruency. (D) Mean accuracy plotted as a function of current trial congruency and volatility. (E) Mean accuracy plotted as a function of proportion congruency and volatility. Circles indicate individual participant values. Error bars indicate standard error. *: p < .05. **: p < .01.

## Results

### Accuracy and response time

Thirty-one participants performed the Stroop task in which they judged the gender of a face while ignoring an overlaid gender word (Figure 1A). Congruency was manipulated by pairing male or female faces with matching or mismatching gender characters, and the proportion congruency varied across stable and volatile blocks (Figure 1B). In volatile runs, the proportion congruency alternated between 20% and 80% across blocks, whereas in stable runs it remained fixed at either 20% or 80%.

We conducted separate 2 (volatility) × 2 (proportion congruency) × 2 (congruency) repeated-measures ANOVAs for reaction time and accuracy. The analyses revealed robust congruency effects on reaction time. The main effect of congruency was significant (F(1, 30) = 185.25, p < 0.001, *η*^2^=0.86), indicating that reaction times were significantly longer under incongruent conditions compared to congruent conditions (Figure 1C). Additionally, there was a significant interaction between stimulus congruency and proportion congruency (F(1, 30) =82.54, p<0.001, *η*^2^=0.73). Post hoc tests revealed that, regardless of the incongruency proportion (20% or 80%), reaction times for incongruent trials were consistently longer than those for congruent trials (20% incongruent proportion: t=13.61, p<0.001, Cohen’s d=2.49; 80% incongruent proportion: t=10.30, p<0.001, Cohen’s d=1.88), but this congruency effect on reaction times was larger in the 20% incongruency condition compared to the 80% condition. Other main effects and interactions related to reaction time were not significant.

For accuracy, the ANOVA revealed a significant two-way interaction between environmental volatility and stimulus congruency (F(1, 30) = 5.53, p = 0.025, *η*^2^=0.16, Figure 1D). In the volatile condition, there was no significant difference in accuracy between the congruent and incongruent trials (t(30) = -0.96, p = 0.34, Cohen’s d= 0.18). Conversely, in the stable condition, accuracy in the incongruent trials was significantly lower than congruent trials (t(30) = -2.62, p = 0.014, Cohen’s d=0.48). The ANOVA also revealed a significant two-way interaction between environmental volatility and proportion congruency (F(1, 30) = 9.85, p = 0.004, *η*^2^=0.25, Figure 1E). In the 20% incongruent blocks, there was no significant difference in accuracy between the volatile and stable conditions (t(30) = -1.65, p = 0.11, Cohen’s d= 0.30). Conversely, in the 80% incongruent blocks, accuracy in the volatile condition was significantly lower than in the stable condition (t(30) = -2.97, p = 0.006, Cohen’s d=0.54). Other main effects and interactions related to accuracy were not significant.

### Model validity analysis

We fit the flexible control model (Figure 2A) to each participant’s reaction speed and observed congruency, separately for each run, yielding trial-wise estimates of learning rate and predicted conflict level. These estimates, together with congruency, were then normalized and multiplied to construct all two-way and three-way interaction terms. Prediction error was subsequently derived as the interaction between predicted conflict level and congruency (–CF × congruency; see Text S1 for details). Using these model-derived variables, we first examined their relation to behavioral performance. Prediction error significantly predicted reaction time, such that larger errors were associated with slower responses (t(30) = 8.84, p < 0.001). By contrast, neither learning rate nor predicted conflict level showed reliable effects on reaction time (learning rate: t(30) = 0.44, p = 0.66; predicted conflict: t(30) = –0.07, p = 0.94). These analyses were performed on orthogonalized variables (see Methods). We then assessed whether the model variables were sensitive to experimental manipulations.

**Figure 2.**
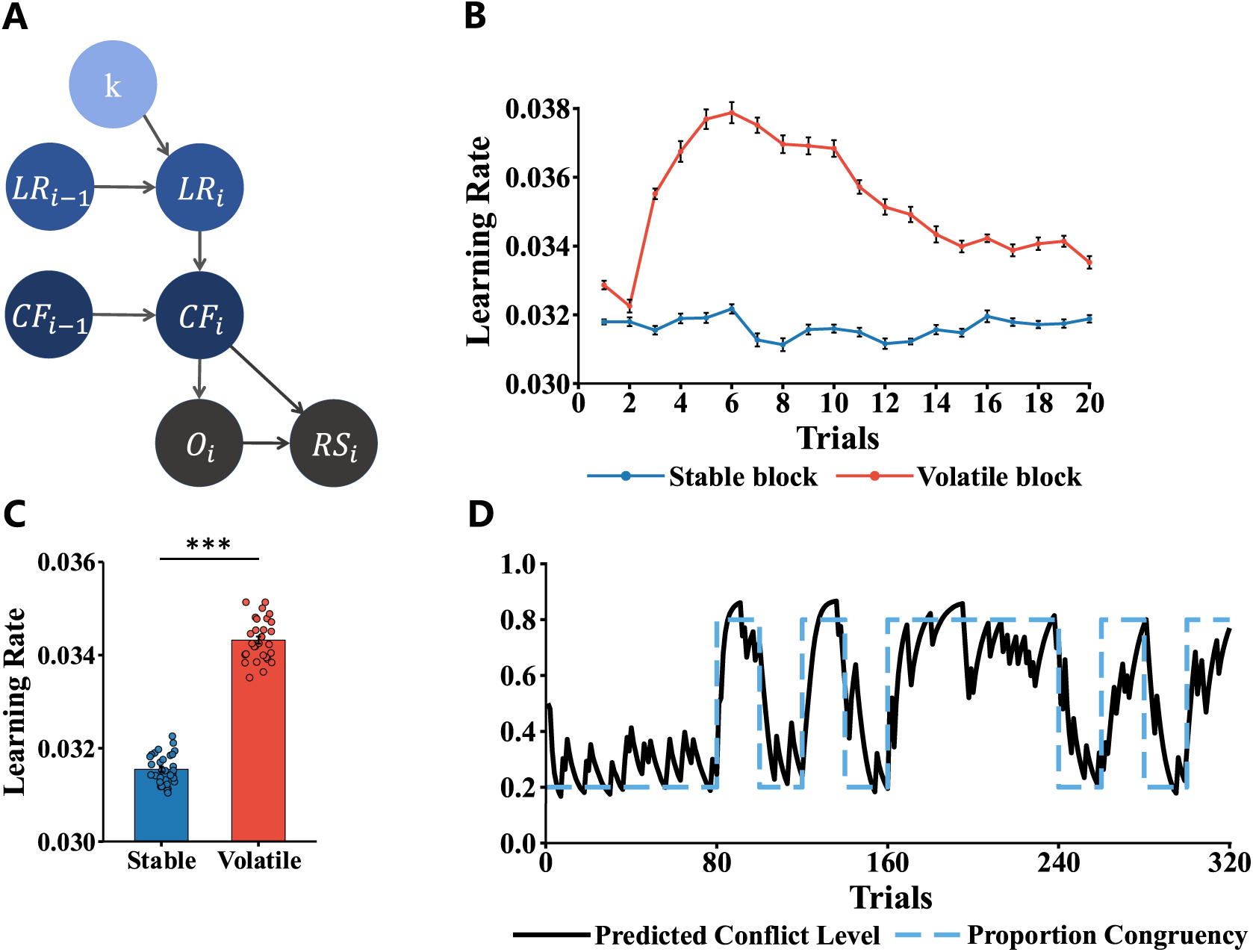
Model structure and validity analysis. (A) Graphical representation of the flexible control model. Model variables are updated on a trial-by-trial basis according to the information flow illustrated in the figure, enabling the system to adapt to control demands of dynamic environment and optimize behavioral performance. Hyperparameter (K): a model hyperparameter held constant within each run, quantifying the probability that the proportion of incongruent trials remains stable during each run. Learning rate (LR): estimate of environmental volatility. Predicted conflict level (CF): the probability that the individual predicts the upcoming trial to be an incongruent trial. Congruency (O): The congruency between the face and the word in the stimulus. Reaction Speed (RS): The speed it takes for a participant to respond to a stimulus. (B) Individual mean model learning rate plotted as a function of run type. (C) The time course of group mean learning rate across trials in volatile and stable blocks. (D) Time courses of the underlying proportion congruency and the corresponding predicted conflict level in an example run.

Learning rate was markedly higher in volatile compared to stable conditions (t(30) = 25.97, p < 0.001; Figure 2C). To capture trial-wise dynamics, we analyzed the 2nd– 4th blocks of each run, averaging learning rate within each trial sequence. In volatile blocks, the learning rate initially increased before gradually decreasing and stabilizing, whereas in stable blocks it remained consistently low (Figure 2B). Finally, driven by the learning rate, predicted conflict levels accurately tracked the switches in proportion congruency throughout the task, despite participants being unaware of when these switches occurred during the experiment (Figure 2D).

### Model comparison

We conducted a model comparison that pitted the flexible control model against the hierarchical Bayesian models with one fixed learning rate (for all runs) or two fixed learning rates (one for stable runs and one for volatile runs)[21]. The models were individually fitted to each of the 31 participants and Bayesian Information Criterion (BIC) values were computed for each participant. When comparing the three models at the group level, the flexible control model demonstrated the highest exceedance probability of 1.0, highlighting its superiority over the other models.

### Relations between single-trial frequency power and model variables

We took the following steps to examine relationships between frequency band power and model variables, and to identify the neural substrates underlying these relationships. We performed these analyses separately for each of the three main model variables (prediction error (PE), learning rate (LR), and predicted conflict rate (CF). In each case we first tested at each electrode whether power in the theta, alpha, or beta band correlated with the model variable across the trials and identified the time–frequency point showing the strongest effect. At this peak, we additionally performed a parallel source localization analysis to determine the underlying brain regions sensitive to the same variable. Both sensor and source level analyses were implemented using linear mixed-effects models. Further methodological details are provided in the Methods section.

### Single-trial Theta and Alpha frequency power related to prediction errors

Analysis using linear mixed-effects models to fit the neural oscillatory power revealed significant correlations of Theta band and Alpha band power by prediction error (PE). In the Theta band, fronto-central electrodes exhibited a positive correlation with PE during the stimulus phase (Figure S1A) with the strongest effect at electrode E80 occurring 534ms after stimulus onset (Figure 3A, t(14210) = 4.71, p < 0.001). Source localization at this peak time point revealed a source in the bilateral medial frontal cortex (MFC) underlying this correlation (Figure 3B, MNI coordinates of peak activation: [10, 50, 40]).

**Figure 3.**
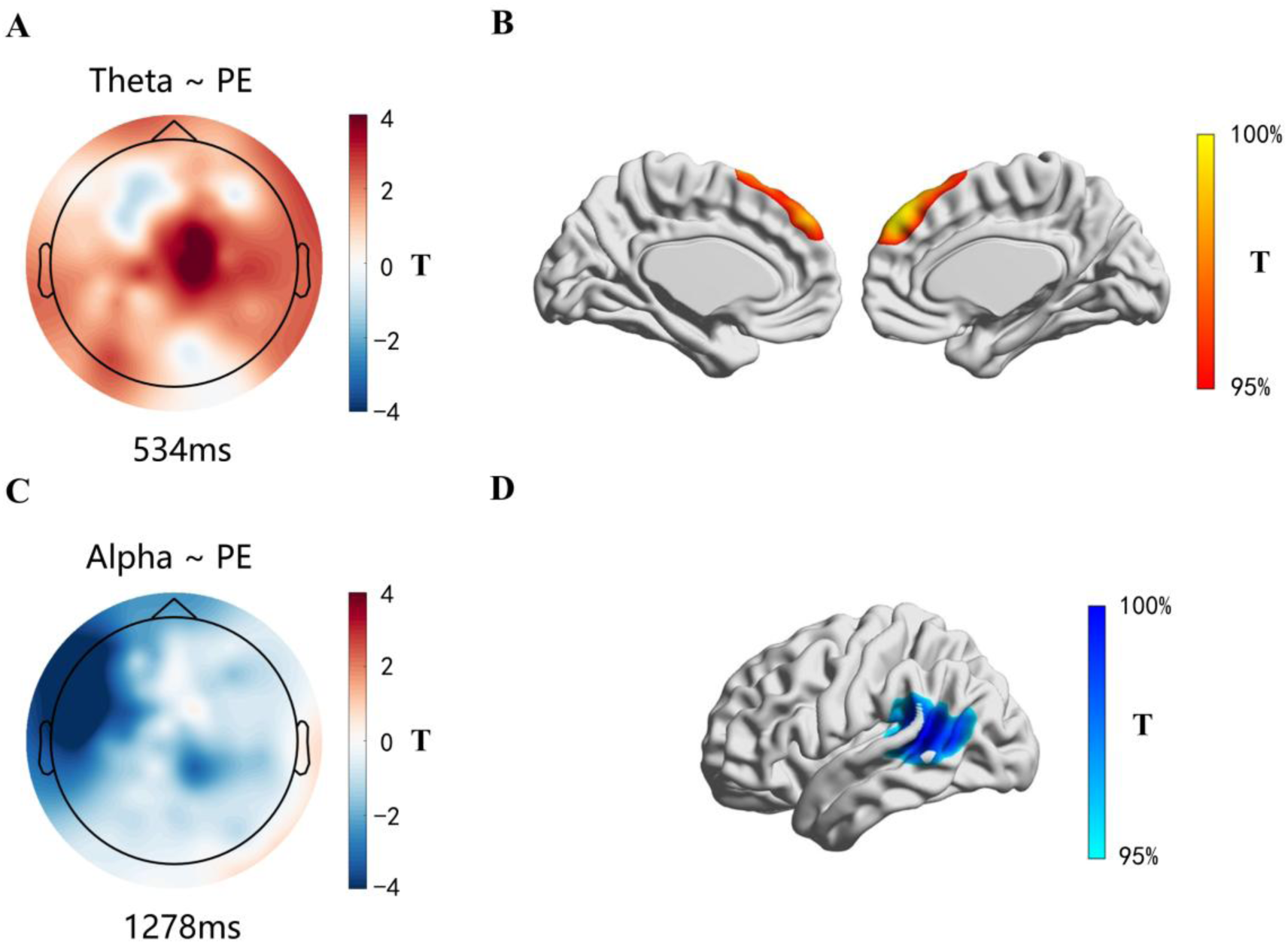
Neural correlations of prediction error (PE). (A, C) Sensor-level topographic maps showing T-value distributions of the correlation between PE and oscillatory power in the Theta (A) and Alpha (C) bands. (B, D) Source-level T-statistic maps corresponding to (A) and (C). In (A), the strongest Theta–PE positive correlation was present over fronto-central electrodes at 534 ms after stimulus onset, with source localization in (B) revealing a top 5% correlation in the bilateral medial frontal cortex. In (C), the strongest Alpha–PE negative correlation was present over left temporal electrodes at 1278 ms, with corresponding source localization in (D) showing a top 5% correlation in the left lateral temporal cortex.

There was also a correlation between the Alpha band power and PE in the left lateral temporal electrodes. This Alpha band correlation extended through both the stimulus phase and the intertrial interval phase (Figure S1B). The strongest correlation was observed at electrode E40, with peak T-value at 1278ms post-stimulus (Figure 3C, t(14210) = -4.86, p < 0.001. Source localization at 1278ms revealed a source in the left temporal cortex, including the superior temporal gyrus, middle temporal gyrus and inferior temporal gyrus (Figure 3D). The peak activation was located in the left middle temporal gyrus (MNI coordinates [-50, -50, 10]).

### Single-trial Alpha and Beta frequency power related to learning rate

Analysis using linear mixed-effects models to fit the neural oscillatory power revealed significant negative correlations of Alpha band and Beta band power by learning rate (LR). As shown in Figure 4A and 4C, significant correlations of learning rate were observed in the right parietal electrodes and left lateral prefrontal electrodes in the Alpha band. These effects lasted from the stimulus onset phase to the end of the intertrial phase (Figure S2A). The strongest correlations by learning rate during the intertrial phase was observed in the right parietal region at electrode E98, 1300ms post-stimulus (figure 4A, t(14210) = -4.22, p < 0.001), and in the left lateral prefrontal region at electrode E34, 1464ms post-stimulus (figure 4C, t(14210) = -5.19, p < 0.001). Further source-level linear mixed-effects model analysis identified the corresponding cortical sources of these correlations at their respective peak time points. The Alpha-LR correlation at 1300ms from the right parietal electrodes corresponded to the right parietal cortex at the source level, including the paracentral lobule, inferior parietal lobule, precuneus, and postcentral gyrus (Figure 4B; peak at the paracentral lobule, MNI coordinates: [10, -30, 79]). The Alpha-LR correlation at 1464ms, corresponding to the left lateral prefrontal electrodes, was primarily localized to the ventrolateral prefrontal cortex (VLPFC), centered on the left inferior frontal gyrus (peak at MNI coordinates: [-30, 30, -22]), and also involved the left insula (Figure 4D).

**Figure 4.**
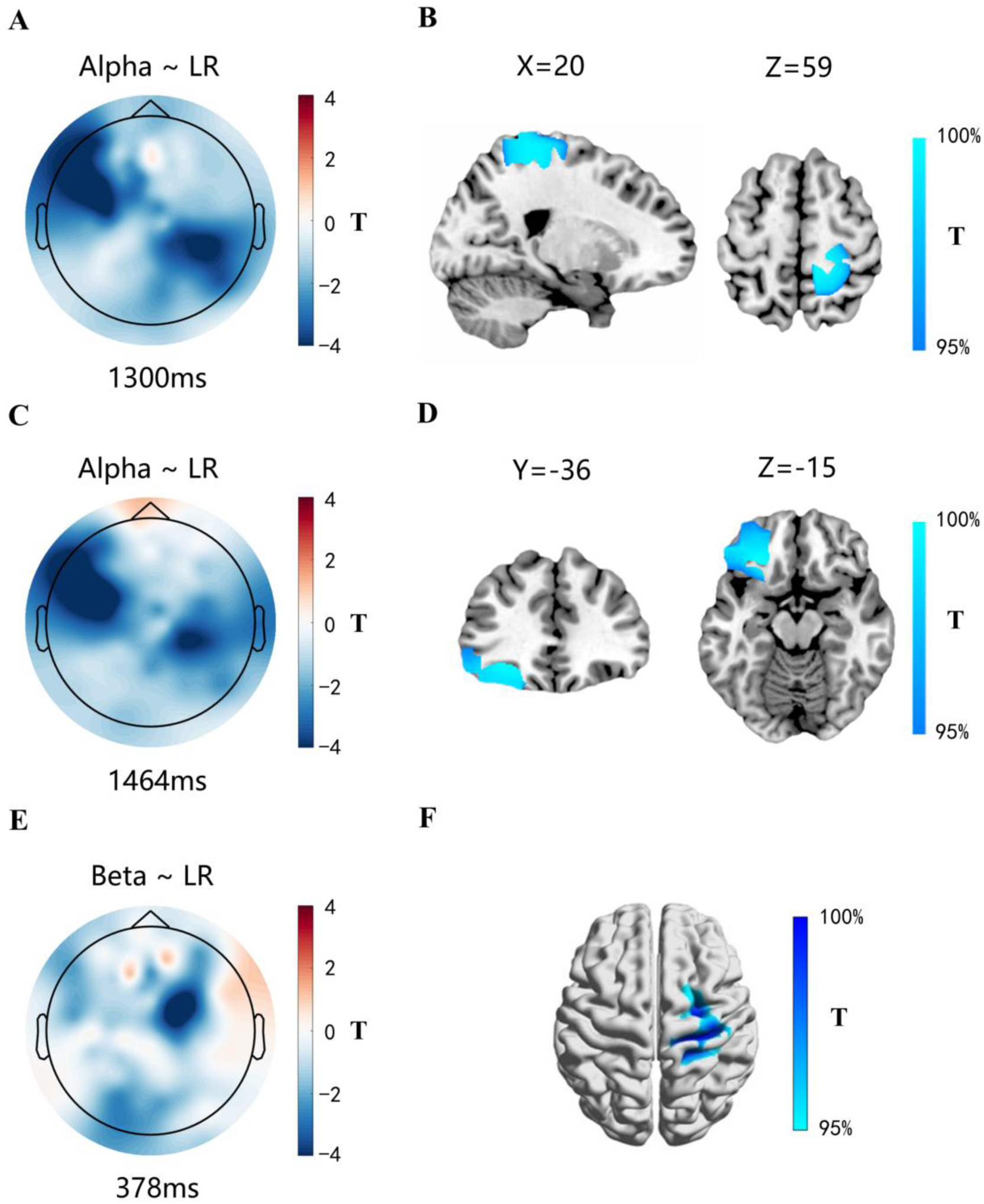
Neural correlations of learning rate (LR). (A, C, E) Sensor-level topographic maps showing T-value distributions of the correlation between LR and oscillatory power in the Alpha (A, C) and beta (E) bands. (B, D, F) Source-level T-statistic maps corresponding to (A), (C), and (E). In (A), the strongest negative Alpha–LR correlation was present over right parietal electrodes at 1300 ms after stimulus onset, with source localization in (B) revealing a top 5% correlation in the right parietal cortex. In (C), the strongest negative Alpha–LR correlation was present over left lateral prefrontal electrodes at 1464 ms, with corresponding source localization in (D) showing a top 5% correlation in the left ventrolateral prefrontal cortex. In (E), the strongest negative Beta–LR correlation was present over right frontal electrodes at 378 ms, with corresponding source localization in (F) showing a top 5% correlation in the right motor cortex.

Learning rate also correlated with Beta band oscillatory power in right frontal electrodes in the stimulus phase (Figure S2B). The strongest correlation was observed at 378 ms post-stimulus at electrode E111 (Figure 4E, t(14210) = -4.97, p < 0.001). Source reconstruction revealed that the Beta-LR correlation at 378 ms was localized to the right precentral and postcentral gyrus at the source level (Figure 4F; peak at the right precentral gyrus, MNI coordinates: [30, -23, 50]).

### Single-trial Alpha frequency power related to predicted conflict level

Analysis using linear mixed-effects models to fit the neural oscillatory power revealed significant positive correlations between Alpha band power and the predicted conflict level that lasted from the stimulus onset phase to the end of intertrial phase (Figure S3). During the intertrial phase, the strongest correlations were found in the right lateral prefrontal region at electrode E111, 1072 ms post-stimulus (Figure 5A, t(14210) = 4.31, p < 0.001), and in the occipitoparietal region at electrode E61, 2016 ms post-stimulus (Figure 5C, t(14210) = 4.09, p < 0.001).

**Figure 5.**
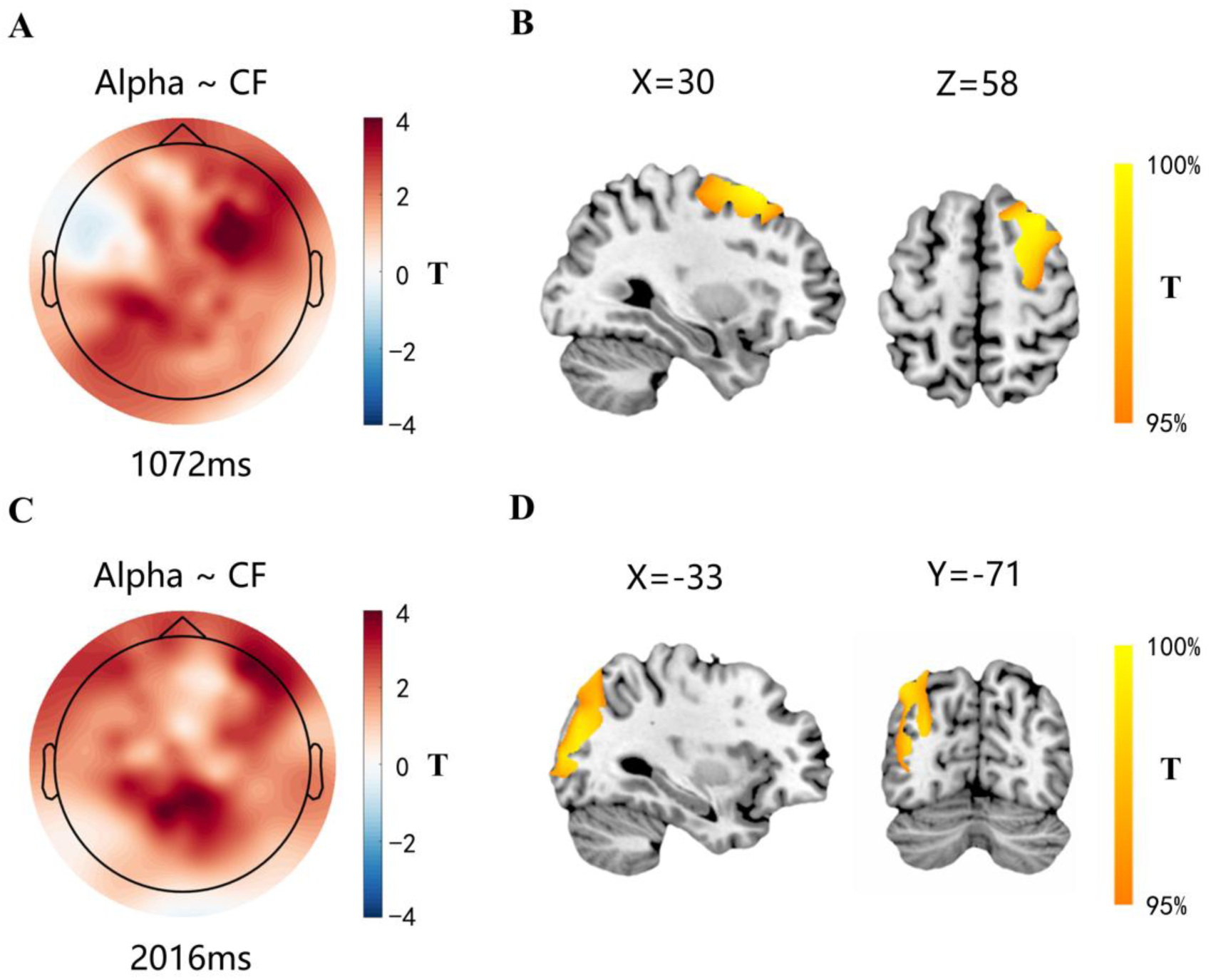
Neural correlations of predicted conflict level (CF). (A, C) Sensor-level topographic maps showing T-value distributions of the correlation between CF and oscillatory power in the Alpha bands. (B, D) Source-level T-statistic maps corresponding to (A) and (C). In (A), the strongest Alpha–CF positive correlation was present over right lateral prefrontal electrodes at 1072 ms after stimulus onset, with source localization in (B) revealing a top 5% correlation in the right dorsolateral prefrontal cortex. In (C), the strongest Alpha–CF positive correlation was present over parieto-occipital electrodes at 2016 ms, with corresponding source localization in (D) showing a top 5% correlation in the left occipitoparietal cortex.

Source localization analysis at the peak time points during the intertrial phase revealed significant neural correlations of predicted conflict level in the right DLPFC including inferior frontal gyrus, middle frontal gyrus and frontal superior gyrus at 1072 ms for the right lateral prefrontal electrodes (Figure 5B, peak at the right frontal superior gyrus, MNI coordinates [30, 20, 63]), as well as in the left occipitoparietal cortex including parietal inferior gyrus, occipital superior gyrus, occipital middle gyrus and occipital inferior gyrus at 2016 ms for the occipitoparietal electrodes (Figure 5D, peak at left occipital middle gyrus, MNI coordinates [-40, -91, 10]).

### Connectivity between frequency band components

The goal of the connectivity analyses was to identify interactions between the frequency band components related to the task parameters that were identified in the preceeding single-trial frequency band analyses. We calculated Granger causality indices reflecting directed information flow between regions encoding model variables during both the stimulus and intertrial phases, and examined directed connections that showed significant increases compare to baseline (see Methods for more details).

As illustrated in Figure 6A, during an early time window(230-640ms) following stimulus onset, the right frontal Beta cluster related to the learning rate exhibited bidirectional information flow with the left lateral prefrontal Alpha cluster related to learning rate, the left temporal Alpha cluster related to prediction error, and the occipitoparietal alpha cluster related to predicted conflict level (see Table S1 and Figure 6A). Within a narrower overlapping time window (420-510ms), shown in Figure 6B, the fronto-central Theta cluster related to prediction error received inflow connections from the right parietal Alpha cluster related to learning rate and the occipitoparietal Alpha cluster related to predicted conflict level. The connections also outflowed from the fronto-central Theta cluster to the left lateral prefrontal Alpha cluster related to learning rate, the left temporal Alpha cluster related to prediction error, and the right lateral prefrontal Alpha cluster related to predicted conflict level (see Table S2 and Figure 6B). The directed information flow between Alpha clusters related to model variables showed a general decrease after stimulus onset, followed by a gradual recovery. During the late intertrial phase, illustrated in Figure 6C, nearly all bidirectional connections showed significantly increased strength relative to baseline, with only two failing to survive correction for multiple comparisons. (see Table S3 and Figure 6C).

**Figure 6.**
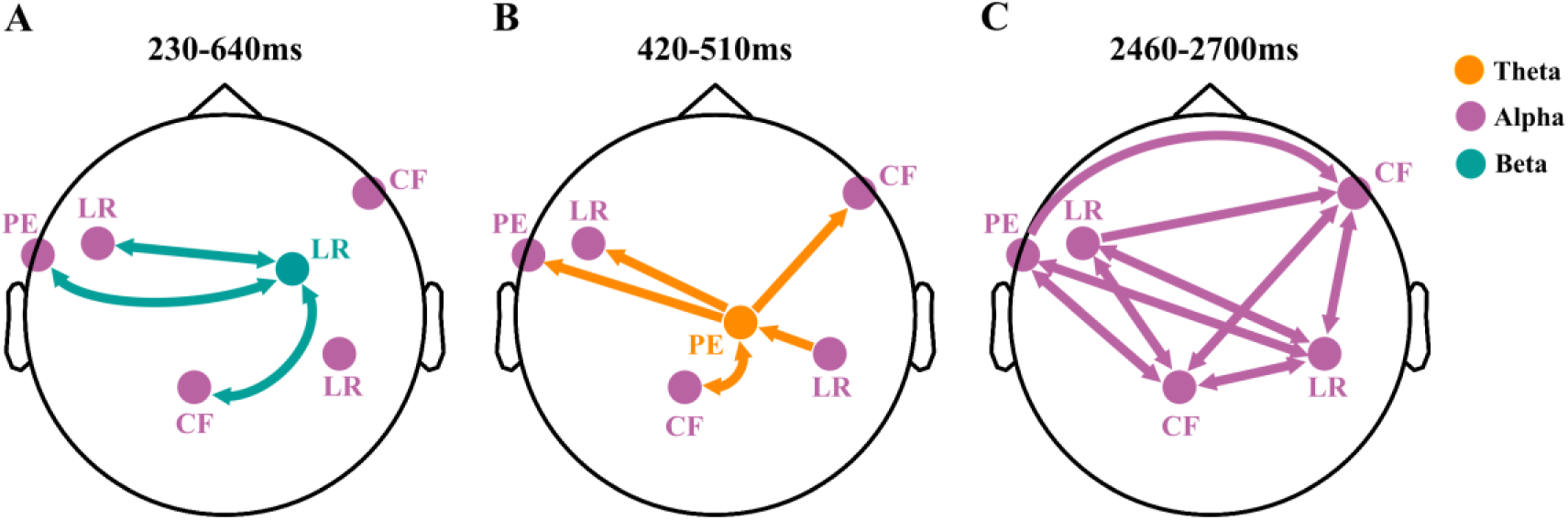
Granger causality connectivity analysis. (A) Directed connections between the Alpha clusters and the Beta cluster during the stimulus phase. (B) Directed connections between the Theta cluster and the Alpha clusters during the stimulus phase. (C) Directed connections between Alpha clusters during the intertrial phase. The time interval indicated at the top of each figure corresponds to the overlap of all time windows in which the connections showed significant increases relative to baseline.

### Connectivity-based correlations of proactive and reactive control

We then related the connectivity patterns revealed by the preceeding Granger causality analyses to an individual measure of control (both proactive and reactive), which we defined as the correlation between prediction error and reaction time. We then identified directed connectivity related to reactive control as that showing a negative modulation by the individual measure of control, and directed connectivity related to proactive control as that showing a positive modulation by the individual measure of control.

Reactive control was triggered by actual conflict to resolve residual conflict quantified by predicted error. This process corresponds to the update of learning rate driven by predicted error in the flexible control model (Figure 7A). We found that the strength of the directed connection from the fronto-central Theta cluster related to prediction error to the left lateral prefrontal Alpha cluster related to learning rate negatively modulated the individual measure of control (average from 330ms to 530ms, R=0.449, P=0.006). and reflects mechanisms involved in reactive control following stimulus onset(Figure 7C).

**Figure 7.**
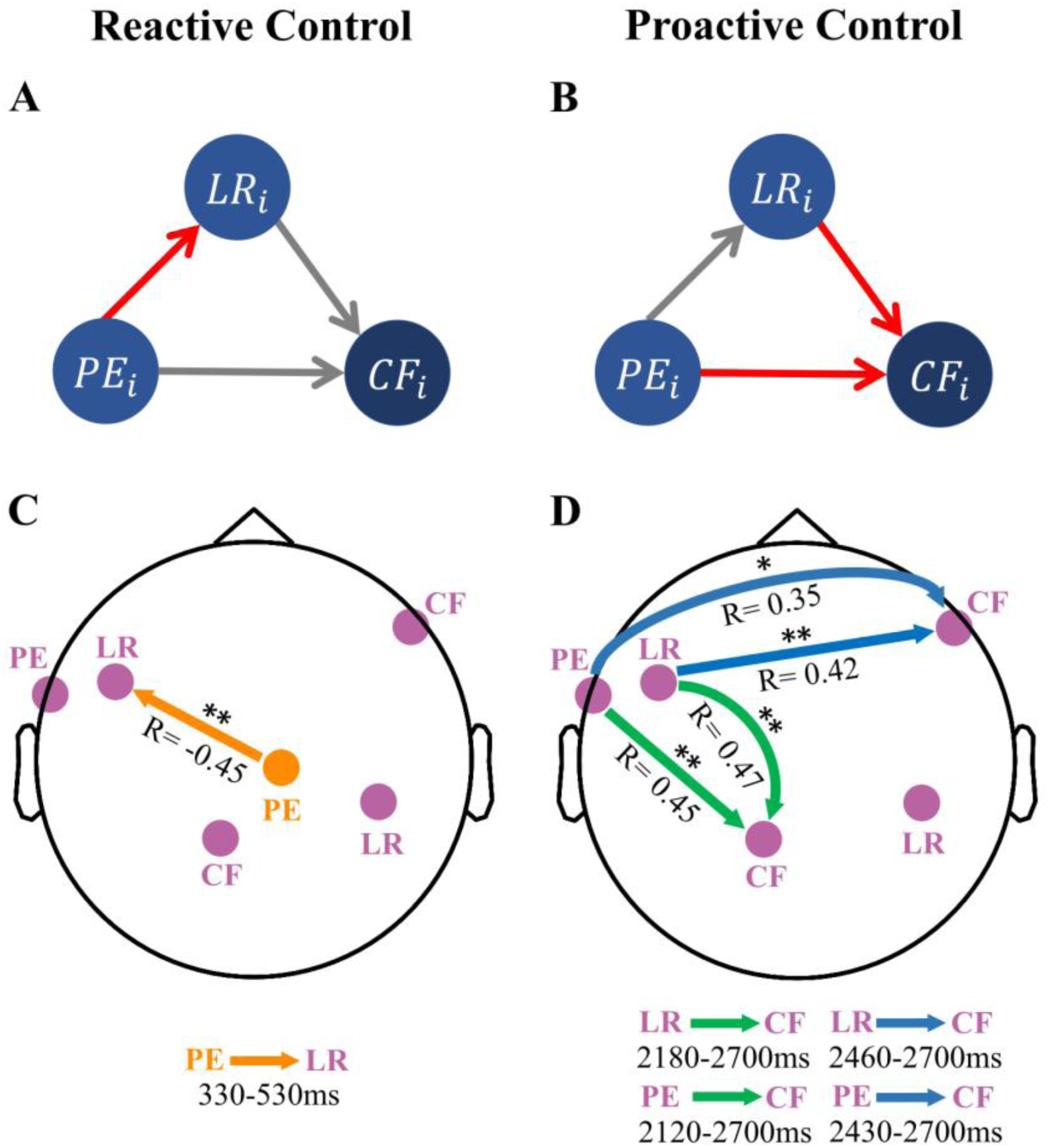
Directional connectivity related to the individual measure of control. (A) and (B) illustrate the flexible control model, highlighting in red the update of learning rate driven by prediction error (A) and the update of predicted conflict level driven by learning rate and prediction error (B), which correspond to reactive and proactive control, respectively. Consistent with the processes characterized in the model, (C) shows that the directional connection from the Theta cluster related to prediction error to the Alpha cluster related to learning rate during the stimulus phase negatively modulated the individual measure of control, providing neural signature of reactive control. (D) shows that directional connections from Alpha clusters related to learning rate and prediction error to Alpha clusters related to predicted conflict level positively modulated the individual measure of control, providing neural signature of proactive control. The legend at the bottom indicates the time intervals over which the average strength of each directed connection was computed; within these intervals, the connection strength was significantly increased relative to baseline. Arrow colors in (C) and (D) indicate the inflow nodes to which the connections were directed, with all other graphical conventions consistent with Figure 6.

Proactive control adjustment corresponds to the update of predicted conflict level, which depends on the information flow from learning rate and prediction error to predicted conflict level in the flexible control model (Figure 7B). We found four Alpha-band directional connections that positively modulated the individual measure of control, and reflected mechanisms involved in proactive control during the late intertrial phase (Figure 7D). Two of these connections inflowed to the occipitoparietal Alpha cluster related to predicted conflict level. One of the two was from the left lateral prefrontal Alpha cluter related to learning rate (connectivity strength average from 2180ms to 2700ms, R=0.468, P=0.004); and the other was from the left temporal Alpha cluster related to prediction error (connectivity strength average from 2120ms to 2700ms, R=0.451, P=0.005). The remaining two directional connections inflowed to right lateral prefrontal Alpha cluster related to predicted conflict level. One outflowed from the left lateral prefrontal Alpha cluster related to learning rate (connectivity strength average from 2460ms to 2700ms, R=0.423, P=0.009) and the other outflowed from left temporal Alpha cluster related to prediction error (connectivity strength average from 2430ms to 2700ms, R=0.349, P=0.027).

## Discussion

We employed a Stroop task in a dynamic environment, together with high-density EEG recording, in order to clarify the temporal process of adaptive adjustments in cognitive control according to environmental changes. We first identified multiple oscillatory frequency band components and corresponding task relevant optimization processes involved in flexible cognitive control. Next, we tested how directed connectivity between these components revealed patterns of information interaction across different task phases. Finally, by relating the strength of specific connections to a behavioral index reflecting individual differences in proactive and reactive control, we obtained evidence that these two modes emerge during distinct phases of the optimization process. Below, we discuss each of these findings in detail. In general, these results provide new insight into the temporal dynamics that link multiple cognitive components to the implementation of adaptive control.

### Relations between single-trial frequency power and model variables

The correlations between model variables and oscillations in different frequency bands reflect distinct cognitive components optimized on a trial-by-trial basis during control. We found that the positive correlation between prediction error and Theta-band oscillations occurred at central–frontal electrodes during the stimulus phase. The Theta-band correlation at these electrodes corresponded to a source in the MFC in the parallel source-level neural correlation analysis. Previous studies have typically regarded the MFC as involved in conflict monitoring, with one of its core functions being the integration and evaluation of control-related information, and its operation being intrinsically linked to Theta oscillations [17,29,31]. The observed correlation between Theta oscillations in this region and prediction error we and others [32] observed provides strong support for the view that conflict monitoring is one of the cognitive components involved in adaptive control.

A negative correlation between prediction error and Alpha-band oscillations occurred at left temporal electrodes during both the stimulus and intertrial phases. The Alpha-band correlation at these electrodes corresponded to a source in the left lateral temporal cortex in the parallel source-level neural correlation analysis. This region overlapped with the superior temporal sulcus (fSTS), which is sensitive to facial detail [40]. Since desynchronization in local Alpha oscillations can enhance corresponding cognitive processing [34,35], increased prediction error may shift cognitive control toward fine-grained processing of facial features, thereby improving gender discrimination accuracy in the subsequent trial. This suggests that prediction error is also involved in feature-based attentional regulation.

Negative correlations between learning rate and Alpha-band oscillations were observed at left lateral prefrontal and right parietal electrodes in both phases. In the parallel source-level neural correlation analysis, the left lateral prefrontal electrodes corresponded to a source in the left vlPFC, and the right parietal electrodes corresponded to a source in the right parietal cortex. Since learning rate quantifies environmental volatility, an increase in learning rate leads to greater uncertainty in the predicted conflict level. In such situations, mental effort pre-allocated based on predicted conflict is more likely to mismatch the actual control demands upon stimulus onset. Therefore, internal task representations need to be reinforced via Alpha-band oscillations related to learning rate to ensure that information processing can not be disrupted even when there is a sudden discrepancy between predicted and actual conflict. Although both the left vlPFC and right parietal cortex are associated with regulating internal attention to resist interference [33,41], there are functional differences between them. The left vlPFC is more involved in resisting interference from dominant semantic features and influencing subsequent adjustment of behavioral responses through cognitive control [42–44]. In contrast, the right parietal cortex is more engaged in maintaining attentional stability by regulating shifts between internal and external attention [41,45]. It is worth noting that previous studies have primarily emphasized the role of the right vlPFC as a component of the ventral attention network [23,44]. However, considering that the lateralized patterns observed across different regions in our study were not restricted to the left vlPFC, we infer that this asymmetry may be related to the type of conflict involved (i.e., stimulus–stimulus conflict in the Stroop task) [46], although prior findings have also reported that the left vlPFC encodes the learning rate [47].

There was also a negative correlation between learning rate and Beta-band oscillations at right frontal electrodes during the stimulus phase, with a source localized to the right motor cortex. In the Stroop paradigm, which primarily involves stimulus–stimulus conflict [46], conflict processing at the response level functions as a necessary complement to conflict processing at the attentional level. This complementary role becomes particularly evident under volatile conditions, where actual conflicts cannot be fully accounted by predicted conflict levels, thereby leading to additional residual conflicts that must be conveyed to the response level for resolution. Such demands require greater motor preparation, reflected in stronger Beta desynchronization [48,49], to ensure efficient response execution. Together, these results indicate that the learning rate is closely involved in both attentional regulation and motor preparation processes.

Positive correlations between predicted conflict level and Alpha-band oscillations were observed at occipitoparietal and right lateral prefrontal electrodes. In the parallel source-level neural correlation analysis, the occipitoparietal electrodes corresponded to a source in the occipitoparietal cortex. Alpha oscillation in the occipitoparietal cortex reflects a gating mechanism by which the brain limits the interference of perceptual information with task-relevant representation [33,50–52]. In the same analysis, the right lateral prefrontal electrodes corresponded to a source in the right DLPFC. Alpha oscillations in the right DLPFC are likely to exert top-down modulation over attentional gating, consistent with its role in modulating downstream information processing based on control demands [53–55].

The relevant cognitive processes in our task include conflict monitoring, attentional selection, and motor preparation. The adjustments of these processes can be interpreted as optimization processes according to the variables of the flexible control model [26]. Our results further demonstrate that changes in oscillatory power across distinct frequency bands underlie these adjustments. However, optimizing these processes in isolation are insufficient to achieve cognitive flexibility. Sustained adaptation to environmental change requires the coordination of these processes, as addressed in the following section.

### Connectivity between frequency band components

Connectivity analyses revealed distinct cross-frequency and within-frequency information flow patterns linking regions associated with the model variables. During the stimulus phase, enhanced Alpha–Beta connectivity prominently involved the right frontal Beta cluster, which was associated with learning rate and localized to the motor cortex. These Alpha–Beta connections combined signals from regions linked to learning rate, predicted conflict, and prediction error, suggesting that both anticipated and unanticipated forms of conflict ultimately required transformation and integration for response. We also observed posterior-to-anterior Alpha–Theta connections with the centro-frontal Theta cluster serving as a relay. These connections conveyed information related to learning rate and predicted conflict; after being integrated and evaluated, these control signals, together with prediction error, provided the basis for optimizing ongoing control processes and were transmitted forward to frontal regions for implementation [17,31]. In the late intertrial phase, a dense pattern of Alpha-band connectivity gradually intensified among regions associated with all three model variables. This information flow enhanced communication between task-relevant regions and likely underlies the updating mechanism of the predicted conflict level, thereby guiding flexible control allocation in preparation for the upcoming conflict, consistent with prior reports linking enhanced Alpha synchronization during preparatory intervals to the facilitation of top-down communication [35]. Taken together, these connectivity patterns illustrate how information flow coordinates distinct frequency components to implement conflict resolution and control allocation processes that adapt to environmental change.

### Connectivity-based correlations of proactive and reactive control

Distinct cognitive components reflected by oscillations in different frequency bands form the basic framework for implementing flexibility in cognitive control, and information communication among these frequency-band components enables the coordination of multiple task-relevant processes. Using the correlation between prediction error and response time as an index of individual differences in proactive and reactive control, we found that, in the late intertrial phase, connections reflecting updates to predicted conflict level showed a positive correlation with this index, indicating that these connections captured individual strength in proactive control. In contrast, in the stimulus phase, connections reflecting learning rate updates showed a negative correlation with this index, indicating that these connections captured individual strength in reactive control.

Our results confirm that the efficiency of predicted conflict updating and learning-rate updating—processes we hypothesized to overlap with proactive and reactive control, respectively—can indeed serve as emergent signatures of these two modes. Taken together, these findings align with the Dual Mechanisms of Control framework, which posits a temporal dissociation between proactive and reactive control (Braver, 2012; Braver et al., 2008). Previous studies have emphasized the link between the MFC, particularly the ACC, and reactive control [21]. We found that individual strength in reactive control is supported by the information flow from the fronto-central Theta cluster to the left lateral prefrontal Alpha cluster, which represents prediction error–driven updates to learning rate. We also observed that individual strength in proactive control is supported by Alpha-band directional connections representing updates to predicted conflict level. The outflow nodes of these connections were the left lateral prefrontal Alpha cluster, related to learning rate, and the left temporal Alpha cluster, related to prediction error. The inflow nodes were the right frontal and the occipitoparietal Alpha clusters, both related to predicted conflict level. Given that the Stroop task primarily engages attentional conflict [46], and that Alpha oscillations underlie attentional selection in our study, it is not surprising that the connectivity patterns capturing both reactive control (Theta–Alpha connectivity) and proactive control (Alpha-band connectivity) prominently involved Alpha oscillations.

## Conclusion

Overall, our study first identified how Theta, Alpha, and Beta oscillations respectively support flexible adjustments within the monitoring, attentional, and motor components of cognitive control in dynamic environments. We further found that the resolution of cognitive conflict is achieved through the coordination of these components via Theta–Alpha and Alpha–Beta directed information flow. Building on this, our analyses revealed that individual differences in proactive and reactive control manifested as emergent signatures in the efficiency of predicted conflict updating during the late intertrial phase and learning-rate updating during the stimulus phase. These findings provide a more precise characterization of the temporal course of adaptive control and the neural architecture that underpins it in dynamic environments, highlighting the critical role of interactions among distinct cognitive components in sustaining cognitive flexibility.

## Methods

### Participants

Thirty-six participants took part in the experiment. All participants were right-handed, had normal or corrected-to-normal vision, and no history of or current neurological impairments. The Ethics Committee of the School of Psychology at South China Normal University approved this study. All participants received monetary compensation upon completion of the experiment. Before the experiment, each participant provided informed consent. Five participants were excluded from further analysis after data acquisition: two participants were excluded due to not following the task instructions, while the other was excluded because of excessive EEG artifacts. In total, the sample size comprised 31 participants (17 female), who were aged from 18 to 27 years old (M = 22.28 ± 2.63 years).

### Materials

We selected 24 neutral facial images (12 male and 12 female) from the Chinese Facial Affective Picture System [56]. Each face image was overlaid with a red Chinese character indicating gender (“男” for male or “女” for female) in SimSun font, size 58. These composite stimuli were displayed against a gray background at the center of the screen. The experiment was programmed using Psychtoolbox extensions in MATLAB 2021a.

### Procedure

Each trial began with the central presentation of a face–character composite stimulus for 1000 ms (Figure 1A). Trials were either congruent (male face with”男”, female face with“女”) or incongruent (male face with”女”, female face with “男”).

Participants were instructed to ignore the overlaid characters and categorize the face gender as quickly and accurately as possible. Responses were made with the left or right index finger using designated keys, with hand-to-gender mapping counterbalanced across participants (e.g., ‘F’ for male with the left hand, ‘J’ for female with the right hand). After stimulus offset, a fixation cross was shown for a variable inter-stimulus interval (2000–3000 ms, steps of 500 ms). To minimize priming effects, no face stimulus was repeated on consecutive trials. Before the main experiment, participants completed a practice session with identical procedures to ensure task comprehension.

The experiment comprised eight runs, each with five blocks. The first block of each run served as a warm-up (24 trials, 50% incongruent) to equalize predictions at run onset. The remaining four blocks contained 20 trials each. Runs were categorized as stable or volatile (Figure 1B). In volatile runs, the proportion of incongruent trials alternated across blocks, shifting from 20% to 80% or vice versa; in stable runs, it remained fixed at either 20% or 80% after the warm-up. The overall number of incongruent trials was matched across stable and volatile runs. The eight runs were divided into two halves, each including two stable and two volatile runs, with run order randomized within each half. Participants were allowed to rest between runs.

### Behavioral data analyses

RT and accuracy were analyzed using a three-way repeated-measures ANOVA with volatility, proportion congruency, and congruency as factors. Error trials, post-error trials, outlier trials, and post-outlier trials were excluded from all RT and EEG analyses. In addition, we fit the flexible control model(see below) to each participant’s trial-wise reaction speed and congruency, separately for each run, yielding trial-wise estimates of learning rate and predicted conflict; estimates corresponding to excluded trials were also discarded. To control for collinearity among variables, the remaining trial-wise estimates were first normalized and then multiplied to construct interaction terms, resulting in seven regressors: learning rate, predicted conflict, congruency, and their two-way and three-way products. Each model variable was subsequently orthogonalized against the others using a general linear model (GLM), and the resulting residuals were used as substitutes for the original variables in subsequent RT and EEG analyses. RT was predicted by each orthogonalized variable within a second GLM, which also included the remaining variables and their products as nuisance predictors to account for shared variance. The slope coefficients of the orthogonalized variables were extracted and tested against zero using one-sample t-tests across participants, allowing us to determine whether each variable significantly modulated RT at the group level.

### Flexible control model

The flexible control model can be described as an extension of the reinforcement learning model [20,21], and its structure originates from decision-making models based on Bayesian networks [57]. The model includes four variables. (1) Learning Rate (LR): Represents environmental volatility and is sensitive to switches in the proportion of incongruent trials. It determines the step size by which the predicted conflict level is updated on each trial. (2) Predicted Conflict Level (CF): Represents the probability that the individual predicts the upcoming trial to be an incongruent trial. (3) Congruency (O): The congruency between the face and the word in the stimulus. (4) Reaction Speed (RS): The speed it takes for a participant to respond to a stimulus, calculated as 1/RT. The variables were updated on a trial-by-trial basis according to the dependency structure illustrated in Figure 2A. In brief, individuals infer learning rate and predicted conflict level according to the hierarchical update rules, and these predictions determine their reaction speeds in subsequent trials associated with stimulus congruency. Therefore, we can infer the expected values of the posterior distributions for the learning rate and predicted conflict level using observed congruency and reaction speeds. Incorporating reaction speed to infer latent variables enables the model to account for individual differences in task performance. We explain the mathematical dependencies among the model variables and how to infer the latent variables based on reaction time and congruency in the Text S1. After model fitting and trialwise normalization of each variable, prediction error (*PE*_*i*_) was defined as −*CF*_*i*_ × *O*_*i*_ for subsequent analysis. The primary variables utilized in subsequent analyses for each participant were learning rate, conflict level, and prediction error.

### Model comparison

We conducted a model comparison that pitted the flexible control model against simpler hierarchical Bayesian models with one learning rate (for the whole task) or two learning rates (one for stable runs and one for volatile runs)[21]. Model performance was quantified using the Bayesian information criterion (BIC).

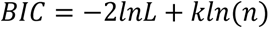

n is the number of trials analyzed, k is the number of free parameters (0, 1, 2 for the flexible control model, the reinforcement learners with one and two learning rates, respectively), and L is the maximized value of the likelihood function. The first term (2*lnL*) can be approximated by *nlnσ*_e_^2^ , where *σ*_e_^2^ is the error variance. The best learning rates for fixed-LR models were selected based on a grid search (from 0.01 to 0.5 with a step size of 0.001) that minimized the BIC. For each subject, the models were fit to the individual trial sequence of congruency and reaction speed, and the simulated reaction speed values derived from each model fit were then used to calculate the BIC, on which the model selection was based. Group-level model comparison was performed using random-effects Bayesian model selection [58] implemented in SPM12.

### EEG acquisition and preprocessing

EEG data were collected using a 129-channel EGI system at a sampling rate of 500 Hz. The Cz electrode served as the reference during recording, and electrode impedances were kept below 50 kΩ. Preprocessing was performed in EEGLAB [59]. The outermost ring of electrodes was excluded because signals at these positions are particularly susceptible to artifacts and their scalp coverage often deviates from standardized electrode coordinates across participants. After exclusion, 102 electrodes remained for analysis. Data were high-pass filtered at 0.1 Hz. For subsequent analyses, different low-pass filters were applied: 35 Hz for the sensor-level neural correlation analysis, and 50 Hz for the source localization and Granger causality analyses. Data were then segmented from –1000ms to 3000ms relative to stimulus onset. Trials with obvious noise were first excluded through manual inspection.

Independent Component Analysis (ICA) was subsequently applied to remove artifacts arising from eye movements, blinks, heart and muscle activity. The cleaned data were re-referenced to the average across the remaining electrodes. Finally, epochs with amplitudes exceeding ±100μV were automatically rejected, resulting in an average of 85.0% ± 10.6% valid trials retained per participant.

### Time–frequency analysis

Time–frequency decomposition was performed in FieldTrip [60] using Morlet wavelet transforms. Analyses focused on Theta (4–7 Hz), Alpha (8–12 Hz), and Beta (21–30 Hz) frequency bands, with wavelet cycle widths of 3, 3, and 7, respectively. The frequency resolution was 0.5 Hz, and the temporal resolution was 2 ms. Because the wavelet window extends beyond the edges of the data segments, values at the beginning and end of each epoch were trimmed, resulting in missing values for the outer time points; thus, the effective analysis time range was shorter than the original segmentation. Power at each time–frequency point was baseline-corrected by subtracting the mean power during a 400–600 ms pre-stimulus baseline and then averaged within each frequency band. From these analyses, single-trial time series of oscillatory power were extracted for subsequent sensor-level neural correlation analyses and cross-frequency Granger causality.

### Model-based neural correlation analysis

We examined the relationship between model variables and oscillatory power using linear mixed-effects models (*fitlme* function in MATLAB). The tested variables were learning rate, predicted conflict level, and prediction error. To minimize collinearity, orthogonalized versions of each variable were used, consistent with the behavioral analyses. For each electrode and sampling point, three separate models were fit for the Theta, Alpha, and Beta bands to test the neural correlation of each variable. In addition to the orthogonalized test variable, all remaining variables and their interactions were included as fixed effects, with participants modeled as random effects. Two linear mixed-effects models were compared—one including only random intercepts, and another including both random intercepts and random slopes. The better-fitting model was chosen based on likelihood ratio tests, and the fixed-effect significance of the test variable in the selected model was used to quantify its neural correlation with oscillatory power.

Single-trial oscillatory power at the source level was further analyzed using the same linear mixed-effects models fitting procedure, but only restricted to the time– frequency points showing the strongest correlation effects with model varilables at the sensor level. For each dipole, t-values of the fixed effects from the selected models were extracted and projected onto the MNI template brain. To identify the oscillatory sources of the sensor-level effects, we focused on the top 5% of voxels showing the strongest neural correlations for each model variable within the same hemisphere corresponding to the sensor-level findings. Anatomical labels for these voxels were determined using the Automated Anatomical Labeling (AAL) atlas, and visualization of t-value distributions was performed with BrainNet [61] and MRIcron.

### Source localization

Dynamic Imaging of Coherent Sources (DICS)[62]; was used to reconstruct single-trial oscillatory power for model-based correlation analyses. A standard anatomical MRI and a boundary element method (BEM) head model from the FieldTrip toolbox were used to construct a 3D template grid at 1 cm resolution in Montreal Neurological Institute (MNI) space. For each frequency band, target time windows were defined based on the strongest neural correlation effects observed at the sensor level, centered on the peak time points and spanning 750 ms (Theta), 500 ms (Alpha), or 350 ms (Beta). A baseline window of equal length was defined, centered 500 ms before stimulus onset. A multitaper frequency transformation was applied to both baseline and target windows to obtain oscillatory power and cross-spectral density. A common spatial filter was computed from the combined baseline and target windows [63] with the regularization parameter set to 10%, and applied to reconstruct single-trial oscillatory power for each frequency band. Power in the target window, baseline-corrected by subtraction of the corresponding baseline power, was then submitted to the linear mixed-effects models fitting procedure described above.

### Granger causality analysis

We performed Granger causality connectivity analysis using the multivariate Granger causality (MVGC) toolbox [64,65]. Data were downsampled to 200 Hz before analysis. Regions of interest (ROIs) were defined as electrode-region pairs correlated with model variables, excluding pairs within the same region. Granger causality was computed both within single frequency bands and across different frequency bands.

For spectral Granger causality within a single frequency band of interest, we first computed time-domain Granger causality, transformed it into the frequency domain, and then averaged the values within the frequency band. For cross-frequency analysis, we obtained the power spectral time series of each band using time–frequency analysis, removed linear trends to improve stability, and then computed Granger causality on the detrended power spectral time series.

For each ROI pair, Granger causality was computed for all electrode pairs within the two ROIs and averaged to represent directed connectivity between regions. Dynamic changes in Granger causality were assessed using a sliding-window approach (500 ms window, 10 ms step) from baseline to the end of the stimulus phase or intertrial phase, depending on the duration of the ROI’s correlation with the model variable.

The optimal model order was determined by calculating the Bayesian Information Criterion (BIC) for autoregressive models at various time lags and recording the optimal order for each model. The median of these optimal orders (15) was then selected as the global model order to ensure consistency across frequency bands, electrode pairs, and time windows [66,67].

Statistical significance for Granger causality was assessed relative to baseline using paired-sample t-tests, with cluster-based permutation testing [68] applied for multiple-comparison correction across the stimulus phase (0–1000 ms), early intertrial phase (1000–2000 ms), and late intertrial phase (2000–2700 ms). Details of baseline selection and cluster-based permutation are provided in Text S2.

### Connectivity-based correlations of proactive and reactive control

We then tested how the connectivity identified in the Granger causalty analyses modulated the individual measure of control, defined on the basis of the correlation between prediction error and response time[21]. A stronger positive correlation reflects a greater reliance on proactive control, because proactive control is deployed based on predicted conflict levels and processes conflict more efficiently when predicted and actual conflict match; whereas a weaker correlation reflects a greater reliance on reactive control, because reactive control can offset the performance impact of residual conflict arising from mismatches between predicted and actual conflict. Since we had attempted to verify in the behavioral data analyses whether prediction error exerts a slowing effect on response times (which was indeed supported, see Results), we further used the t-values from the same GLM, which reflected how orthogonalized prediction error explained variance in response times and served as the indicator of this correlation. For reactive control, we tested whether the strength of directed connections from regions related to prediction error to those related to the learning rate during the stimulus phase (as identified in the connectivity analysis) negatively modulated the correlation between prediction error and response time, as these directed connections may provide neural signature for reactive control (Figure 7A). For proactive control we tested whether the strength of directed connections from regions related to the learning rate or prediction error to regions related to the predicted conflict level during the late intertrial phase(as identified in the connectivity analysis) positively modulated the correlation between prediction error and response time, as these directed connections may provide neural signature for proactive control (Figure 7B).

This method has the potential to identify multiple directed connections that potentially support proactive control during the intertrial phase and reactive control during the stimulus phase. For each connection, we averaged the connection strength over the time intervals during which it was significantly increased relative to baseline. To assess their relationship with these two control modes, we computed the correlation between these connection strengths and the T-values reflecting the effect of prediction error on response time. To control for multiple comparisons, the significance of these correlations was FDR-corrected separately for potential connections related to proactive and reactive control according to the number of tests in each set.

## Acknowledgement

This research was supported by the National Natural Science Foundation of China (Grant No. 32571283) and the National Science and Technology Innovation 2030 Major Program (Grant No. 2021ZD0203800).

