## Supplementary Information for "Temporal Dynamics of Flexible Cognitive Control"

This Supplementary Information includes:

Text S1

Text S2

Figure S1

Figure S2

Figure S3

Table S1

Table S2

Table S3

**Text S1. Flexible control model**

This section describes the conditional distributions that define dependencies among variables, and the posterior inference procedure used to compute their trialwise values. For additional details, see Jiang and colleagues [20,21].

For the $i+1$-th trial, its learning rate $LR_{i+1}$ has a probability $k$ of remaining the same as $a_i$, and a probability $1-k$ of randomly jumping to any value within its range (0 to 1):

$$p(LR_{i+1}|LR_i, k) = 1 - k + k\delta(LR_{i+1} - LR_i) \tag{1}$$

Where $\delta$ is the Dirac delta function. Since participants in our task were not explicitly informed about the switching rules of the proportion of incongruent trials, this setup reflects the fact that, regardless of whether in volatile or stable conditions, the probability that the next trial is incongruent will, in most cases, remain consistent with the current trial. By using a uniform distribution to model the changes in the learning rate, we avoid introducing excessive prior assumptions that might constrain its variability. The results depicted in Figure 2B demonstrate that the posterior estimates of the learning rate are sensitive to the underlying changes in the proportion of congruency. In volatile condition blocks, the learning rate shows an initial increase followed by a gradual leveling off, indicating adaptation to changing control demands. In contrast, in stable condition blocks, the learning rate remains at a relatively low level throughout (averaged over blocks 2–4 for runs in both volatile and stable conditions), reflecting the consistent environment.

Given the randomness of congruency sequence, it is impossible to make a precise prediction of conflict. Hence, the prediction should be approximate, leading to a smooth distribution of the predicted conflict level. In our model, the uncertainty of the prediction depends on the learning rate $a_{i+1}$. To mathematically describe this relationship, we introduce the variable volatility $v_{i+1}$, defined as:

$$v_{i+1} = \frac{1}{LR_{i+1}} - 2 \quad (2)$$

Subsequently, we model the intermediate predicted conflict level $CF_{i+0.5}$ using a Beta distribution:

$$CF_{i+0.5} \sim Beta(CF_i v_{i+1} + 1, v_{i+1} - CF_i v_{i+1} + 1) \quad (3)$$

In this distribution, the mode is $CF_i$, and the sum of the two shape parameters is $\frac{1}{LR_{i+1}}$. The reason for using the Beta distribution to define the uncertainty of the predicted conflict level is that it can be interpreted as the likelihood function of observing $CF_i v_{i+1}$ incongruent trials and $v_{i+1} - CF_i v_{i+1}$ congruent trials. According to this interpretation, $v_{i+1}$ reflects the number of past trials considered by the individual when updating predicted conflict level. Finally, we update the predicted conflict level using the form of a standard reinforcement learning rule:

$$CF_{i+1} = CF_{i+0.5} + LR_{i+1}(O_i - CF_{i+0.5}) \quad (4)$$

This update adjusts the predicted conflict level $CF_{i+1}$ based on the discrepancy between the observed congruency $O_i$ and the intermediate predicted conflict level $CF_{i+0.5}$, scaled by the learning rate $LR_{i+1}$.

After the occurrence of the next observed congruency, the individual's reaction speed (where $RS_i$=1/$RT_i$, $RS_i$ was used because of its normality) follows an approximately Gaussian distribution conditioned on the predicted conflict level $CF_{i+1}$

and the observed congruency $O_{i+1}$:

$$P(RS_{i+1}|O_{i+1}, CF_{i+1}) \propto (1 - |O_{i+1} - CF_{i+1}|)e^{-(RS_{i+1} - a_{o_{i+1}} CF_{i+1} - b_{o_{i+1}})^2 / 2\sigma_{o_{i+1}}^2} \quad (5)$$

Where $|O_{i+1} - CF_{i+1}|$ quantified the discrepancy between estimated and actual conflict. $a_{o_{i+1}}$, $b_{o_{i+1}}$, and $\sigma_{o_{i+1}}$ are congruency-specific hyperparameters to be estimated. Since $O_{i+1}$ can be either 0 (congruent trial) or 1 (incongruent trial), there is a separate set of hyperparameters $a_{o_{i+1}}$, $b_{o_{i+1}}$, and $\sigma_{o_{i+1}}$ for each congruency type. These hyperparameters were optimized via the expectation-maximization (EM) algorithm. Specifically, we modeled reaction speed as a function of predicted conflict level using a General Linear Model (GLM). For the two congruency types, we obtained separate sets of parameters $a_{o_{i+1}}$, $b_{o_{i+1}}$, and $\sigma_{o_{i+1}}$. where $a_{o_{i+1}}$ represents the slope, $b_{o_{i+1}}$ is the intercept, and $\sigma_{o_{i+1}}$ denotes the residual standard deviation of the GLM fit. Then, we replaced the old parameter sets with the new ones and re-estimated the predicted conflict levels using these updated parameters. This updated set of predicted conflict levels was then used to fit the reaction speeds again, generating further updates to the hyperparameters. This iterative process continued until the parameter sets converged.

To update the estimate of internal variables (predicted conflict level $CF_{i+1}$ and learning rate $LR_{i+1}$) based on the reaction speeds and observed congruency, we employed a posterior inference process on a trial-by-trial basis. The estimates of learning rate and predicted conflict level were computed as the expectation of their corresponding marginalized distributions:

$$P(k, LR_{i+1}, CF_{i+1} | O_1, RS_1 \dots\dots O_i, RS_i) \propto \iint P(k, LR_i, CF_i | O_1, RS_1 \dots\dots O_{i-1}, RS_{i-1}) \times$$

$$P(RS_i | O_i, CF_i) \times P(LR_{i+1} | LR_i) \times P(CF_{i+1} | LR_{i+1}, O_i) dLR_i dCF_i \quad (6)$$

Since all model variables were standardized for subsequent analyses, prediction error (PE) was also redefined accordingly:：

$$PE_i = 1 - O_i CF_i \quad (7)$$

where both $O_i$, $CF_i$ were normalized. Because the normalized congruency $O_i$ only takes values of -1 or 1, we further scaled $CF_i$ to the range of [-1, 1]. However, given that the current analyses focus on the linear relationships between model variables and either reaction time or neural oscillations, scaling $CF_i$ is not strictly necessary. Therefore, Equation (7) can be further simplified as：

$$PE_i \propto -O_i CF_i \quad (8)$$

**Text S2. Cluster-based permutation for Granger causality analysis**

For spectral Granger causality within a single frequency band, the baseline period matched that used in the time–frequency analysis (−600 to −400ms). For cross-frequency Granger causality, however, using the same baseline was not feasible because both time–frequency decomposition and Granger estimation apply sliding windows, resulting in the baseline being trimmed twice. To avoid this and ensure the baseline was free from stimulus-evoked activity, we used −350 to −250ms as the baseline for cross-frequency analysis.

Significance during the post-stimulus period was assessed relative to baseline using paired-sample t-tests, with cluster-based permutation testing [68] applied for multiple-comparison correction. Because our subsequent analyses aimed to test the temporal dissociation hypothesis of proactive and reactive control within the set of previously identified connections, and given that prior studies typically place proactive control deployment before next trials while regarding reactive control as stimulus-triggered [2], we performed corrections separately within the broad analysis epoch (0–2700ms) for the stimulus phase (0–1000ms), early intertrial phase (1000–2000ms), and late intertrial phase (2000–2700ms). The alpha level for single-time comparisons was set to 0.05, and the cluster-level alpha to 0.05 (two-sided), to identify time intervals showing significant changes in directional connectivity.

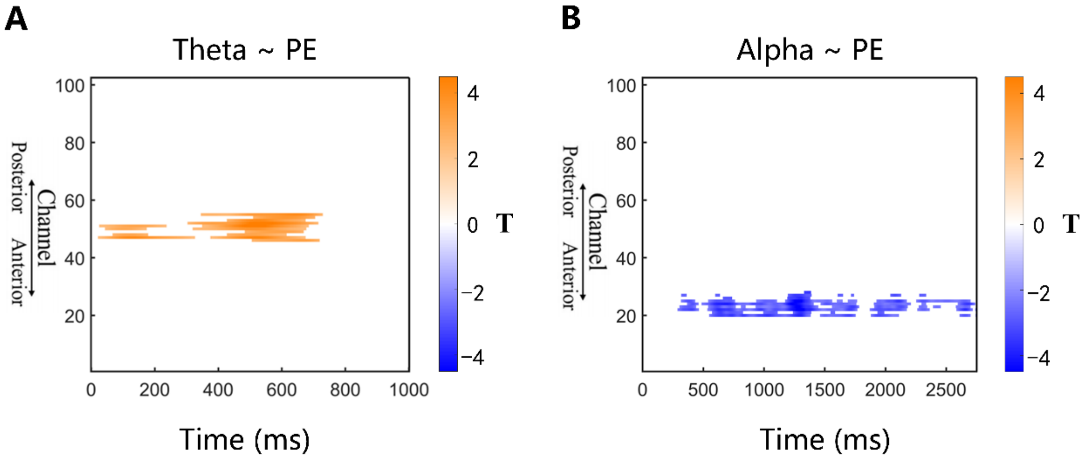


**Figure S1. Neural correlates of prediction error (PE) for each time point.** (A) Theta band power was significantly related to trial-wise predicted error during the stimulus phase. (B) Alpha band power was significantly related to trial-wise predicted error during both the stimulus and intertrial interval phases.

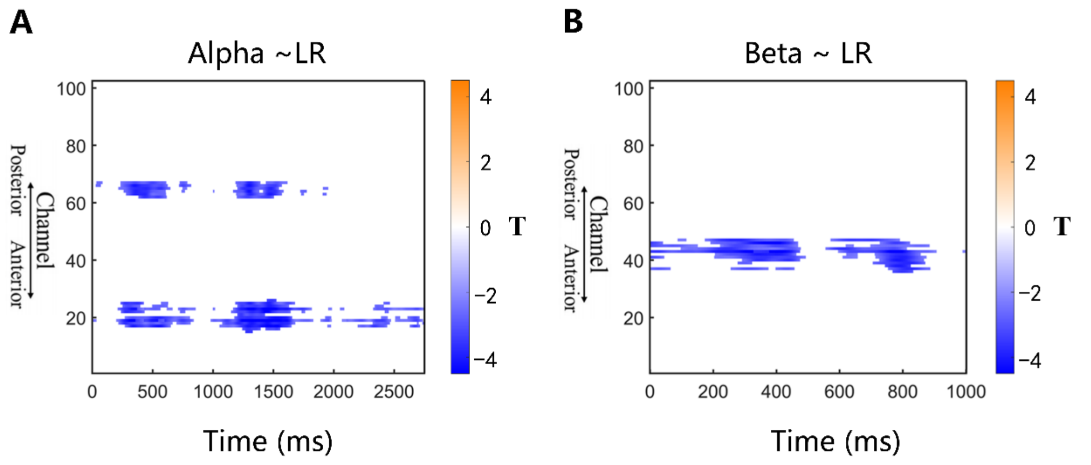


**Figure S2. Neural correlates of learning rate (LR) for each time point.** (A) Alpha band power was significantly related to trial-wise learning rate during both the stimulus and intertrial interval phases. (B) Beta band power was significantly related to trial-wise learning rate during the stimulus phase.

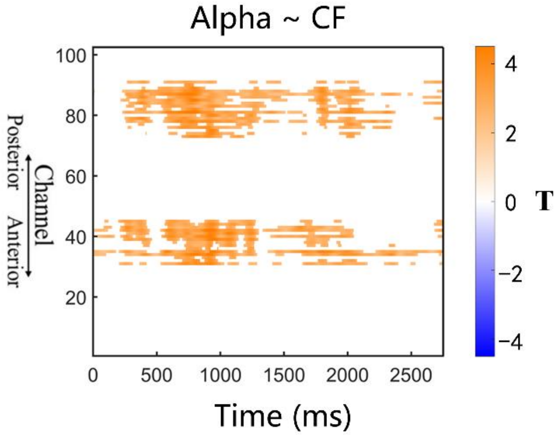


**Figure S3. Neural correlates of predicted conflict level (CF) for each time point.**

Alpha band power was significantly related to trial-wise predicted conflict level during both the stimulus and intertrial interval phases.

**Table S1. Alpha-Beta cross frequency increased directional connectivity in the stimulus phase.**

| Inflow node(frequency band/related variable) | Outflow node(frequency band/related variable) | Time interval(ms) |
|---|---|---|
| Left Lateral Prefrontal(Alpha/LR) | Right frontal(Beta/LR) | 230-640 |
| Right frontal(Beta/LR) | Left Lateral Prefrontal(Alpha/LR) | 100-700 |
| Left Temporal(Alpha/PE) | Right frontal(Beta/LR) | 220-640 |
| Right frontal(Beta/LR) | Left Temporal(Alpha/PE) | 190-700 |
| Occipitoparietal(Alpha/CF) | Right frontal(Beta/LR) | 140-1000 |
| Right frontal(Beta/LR) | Occipitoparietal(Alpha/CF) | 160-870 |

**Table S2. Theta-Alpha cross frequency increased directional connectivity in the stimulus phase.**

| Inflow node(frequency band/related variable) | Outflow node(frequency band/related variable) | Time interval(ms) |
|---|---|---|
| Left Lateral Prefrontal(Alpha/LR) | Frontal-central(Theta/PE) | 330-530 |
| Left Temporal(Alpha/PE) | Frontal-central(Theta/PE) | 340-520 |
| Frontal-central(Theta/PE) | Right Parietal(Alpha/LR) | 420-590 |
| Right Lateral Prefrontal(Alpha/CF) | Frontal-central(Theta/PE) | 340-510 |
| Frontal-central(Theta/PE) | Occipitoparietal(Alpha/CF) | 250-570 |
| Occipitoparietal(Alpha/CF) | Frontal-central(Theta/PE) | 340-550 |

**Table S3. Alpha band increased directional connectivity in the late intertrial phase.**

| Inflow node(related variable) | Outflow node(related variable) | Time interval(ms) |
|---|---|---|
| Left Temporal(PE) | Right Parietal(LR) | 2330-2700 |
| Left Temporal(PE) | Occipitoparietal (CF) | 2310-2700 |
| Right Parietal(LR) | Occipitoparietal (CF) | 2350-2700 |
| Right Parietal(LR) | Left Temporal(PE) | 2240-2700 |
| Right Parietal(LR) | Left Lateral Prefrontal(LR) | 2280-2700 |
| Right Parietal(LR) | Right Lateral Prefrontal(CF) | 2560-2700 |
| Left Lateral Prefrontal(LR) | Right Parietal(LR) | 2430-2700 |
| Left Lateral Prefrontal(LR) | Occipitoparietal(CF) | 2220-2700 |
| Occipitoparietal(CF) | Left Lateral Prefrontal(LR) | 2180-2700 |
| Occipitoparietal(CF) | Right Parietal(LR) | 2190-2700 |
| Occipitoparietal(CF) | Right Lateral Prefrontal(CF) | 2230-2700 |
| Occipitoparietal(CF) | Left Temporal(PE) | 2120-2700 |
| Right Lateral Prefrontal(CF) | Right Parietal(LR) | 2280-2700 |
| Right Lateral Prefrontal(CF) | Occipitoparietal(CF) | 2120-2700 |

| | | |
|---|---|---|
| Right Lateral Prefrontal(CF) | Left Lateral Prefrontal(LR) | 2460-2700 |
| Right Lateral Prefrontal(CF) | Left Temporal(PE) | 2430-2700 |
